# Genetic drivers of repeat expansion disorders localize to 3-D chromatin domain boundaries

**DOI:** 10.1101/191213

**Authors:** James Sun, Linda Zhou, Daniel J. Emerson, Thomas G. Gilgenast, Katelyn Titus, Jonathan A. Beagan, Jennifer E. Phillips-Cremins

## Abstract

More than 25 inherited neurological disorders are caused by the unstable expansion of repetitive DNA sequences termed short tandem repeats (STRs). A fundamental unresolved question is why specific STRs are susceptible to unstable expansion leading to severe pathology, whereas tens of thousands of normal-length repeat tracts across the human genome are relatively stable. Here, we unexpectedly discover that nearly all STRs associated with repeat expansion diseases are located at boundaries demarcating 3-D chromatin domains. We find that boundaries exhibit markedly higher CpG island density compared to loci internal to domains. Importantly, disease-associated STRs are specifically localized to ultra-dense CpG island-rich boundaries, suggesting that these loci might be hotspots for epigenetic instability and topological disruption upon unstable expansion. In Fragile X Syndrome, mutation-length expansion at the *Fmr1* gene results in severe disruption of the boundary between TADs. Our data uncover higher-order chromatin architecture as a new dimension in understanding the mechanistic basis of repeat expansion disorders.

---

Unstable expansion of short tandem repeats (STRs) serves as the mechanistic basis for more than 25 inherited neurological disorders, including Fragile X syndrome, Huntington’s disease, Amyotrophic lateral sclerosis and Friedreich’s ataxia ^1–5^. Disease-associated STR tracts (hereafter referred to as daSTRs) exhibit tremendous diversity in sequence, gene body location, and tract length, rendering them challenging to study and raising the question of whether an integrative model could exist to explain repeat instability. Healthy individuals have tens of thousands of normal-length STRs distributed throughout their genomes ^6–8^. Normal-length STR tracts are generally stable across generations and among somatic tissues in the same individual ^4^. By a process that is poorly understood, some critical normal-length alleles undergo somatic or germ line expansion and transition to intermediate, pre-mutation and mutation (affected) repeat unit tract lengths ^1,5^, leading to disease. A fundamental unresolved question is why some key genes are susceptible to unstable expansion leading to severe pathology, whereas many genes appear to tolerate and stably propagate normal-length STRs.

Mammalian genomes are organized into a complex hierarchy of highly self-interacting Megabase (Mb)-scale structures termed topologically associating domains (TADs) ^9,10^ and smaller, nested subTADs ^9–12^. TADs/subTADs span >90% of the genome and are thought to create insulated neighborhoods demarcating the search space of specific long-range interactions between enhancers and their target genes ^13–15^. To shed new light on the difference between normal-length repeats known to undergo unstable expansion and those that remain stable, we first explored higher-order 3D genome folding patterns around daSTR loci in Hi-C maps generated from human embryonic stem (ES) cells ^16^ and cortical tissue ^17^. Because the samples are not diseased, we mapped daSTRs to the normal-length repeat tracts found in the hg19 reference genome (**Supplementary Table 1**). To quantitatively determine the precise location of TAD boundaries, we first explored 3,062 TADs reported in pluripotent human ES cells with a well-established method based on a directionality index (DI) test statistic and hidden Markov model (DI+HMM) ^9,16^ (**Supplementary Fig. 1A**). Surprisingly, we observed that 15 out of 27 daSTR loci exhibit striking co-localization with boundaries of TADs, including *FMR1* (fragile X syndrome), *HTT* (Huntington’s disease), *DMPK* (Myotonic Dystrophy 1), *FXN* (Friedreich’s Ataxia), *C9orf72* (Amyotrophic Lateral Sclerosis) and *ATXN1* (spinocerebellar ataxia 1) (**Fig. 1A, Supplementary Fig. 2**).

**Figure 1.**
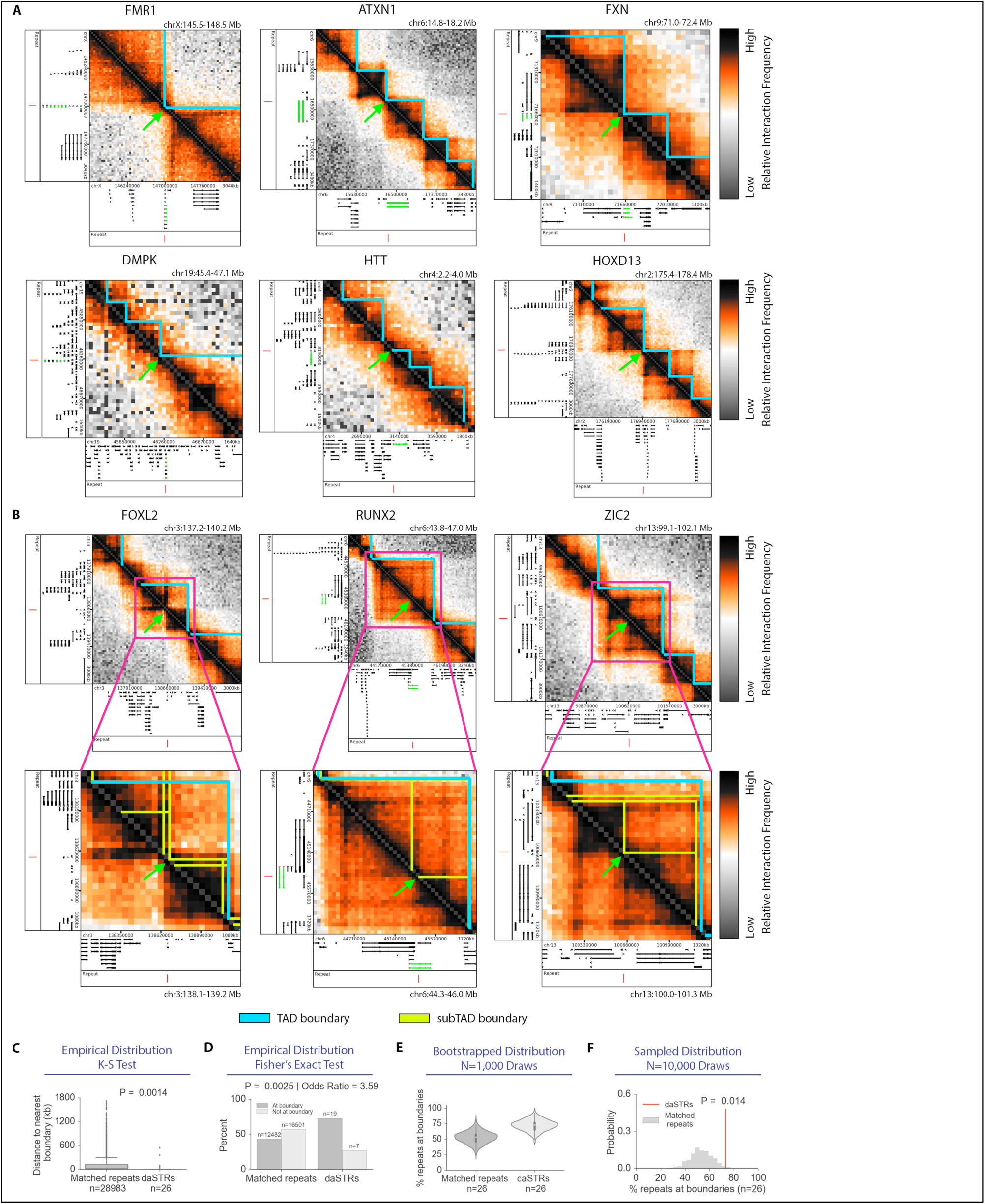
Nearly all disease-associated short tandem repeats (daSTRs) susceptible to pathologic instability are localized to 3D chromatin domain boundaries. (**A-B**) Heatmaps of 40 kilobase binned Hi-C data in human embryonic stem cells. **(A)** daSTR loci co-localized with TAD (blue) and **(B)** subTAD (yellow) boundaries. Genes (green) containing the daSTR (red) are shown in the tracks below heatmaps. Domain boundaries at daSTRs are demarcated with a green arrow. **(C)** Empirical distribution of genomic distance from daSTRs and matched repeats to the nearest domain boundary. Of the 27 daSTRs analyzed in this study, CSTB was dropped from the statistical test because normal-length matched repeats were not found in the hg19 reference genome. **(D)** Bar plots comparing placement of daSTRs and matched repeats at boundaries or internal to domains. **(E)** Bootstrapped distributions of percent daSTRs or matched repeats overlapping boundaries. **(F)** Percent daSTRs overlapping boundaries compared to an empirical null distribution consisting of the percent matched, normal-length repeats.

To better understand the daSTRs initially classified as distal from TAD boundaries, we identified a nested hierarchy of 12,274 TAD and subTADs in human ES cells (**Supplementary Fig. 1B**) by performing a sweep of the DI parameter as detailed in the **Supplementary Methods**. We identified an additional 5 out of 27 daSTRs at subTAD boundaries, including *RUNX2* (cleidocranial dysplasia), *ZIC2* (holoprosencephaly) and *CACNA1A* (spinocerebellar ataxia 6) (**Fig. 1B, Supplementary Fig. 3**). Notably, 5 of the 7 daSTR loci that did not exhibit adjacency to quantitatively called domain boundaries were still co-localized at visually evident domain borders (**Supplementary Figs. 4-5**). We attribute the false negative boundary calls to limitations in the sensitivity of the DI method in calling nested domains, suggesting that our identified 20 out of 27 daSTR loci at human ES cell boundaries is a conservative estimate. Together, our analyses indicate that nearly all daSTRs susceptible to unstable expansion are spatially proximal to boundaries of 3D genome domains in human ES cells **(Supplementary Fig. 6)**.

We next set out to determine how the boundary localization of daSTRs compared to the genome-wide expectation of matched normal-length repeats. To ensure a rigorous null model, we compared the daSTR loci in hg19 to all other normal-length STR tracts (3-32 repeat units) matched by repeat sequence and gene body placement (**Supplementary Methods**). We found that daSTRs are significantly closer to domain boundaries compared to matched repeat tracts (0 bp versus 19,325 bp median distance; P = 9.93e-7 Mann Whitney U Test; P = 0.0014, Kolmogorov-Smirnov test) (**Fig. 1C)**. Moreover, daSTRs showed significantly higher enrichment at domain boundaries compared to other normal-length, matched repeat sequences (Odds Ratio = 3.59, Fisher’s Exact Test, P = 0.0025, **Fig. 1D**). This striking boundary enrichment is a conservative estimate given the high number of daSTRs located at visually apparent domain boundaries missed by the DI+HMM method. To account for the possibility of false positives/negatives in the DI+HMM domain detection method, we computed bootstrapped confidence intervals for the percentage of repeat tracts located at boundaries (**Supplementary Methods**). The mean percentage of drawn repeats tracts that are boundary-associated increased from 53.5% (bootstrapped 95% CI: 34.5% < μ_percent_boundary_ < 72.5%) in matched repeats compared to 73.1% (bootstrapped 95% CI: 55.9% < μ_percent_boundary_ < 90.3%) in daSTRs, respectively (**Fig. 1E**). Finally, we conducted a randomization test and demonstrated that daSTRs are significantly closer to domain boundaries compared to the genome-wide null distribution of matched repeats (empirical P=0.018, **Fig. 1F**). Together, our data indicate that loci susceptible to pathologic, unstable repeat expansion are significantly closer to domain boundaries than expected genome-wide by matched, normal-length repeat sequences. Our findings are robust across several empirical and non-parametric statistical tests.

We next set out to determine if the strong enrichment of daSTRs at domain boundaries was specific to ES cells or more generalizable across lineages and species. Due to the read depth and resolution limits of Hi-C data published to date, subTAD boundaries have not been reported genome-wide across multiple human cell types. Therefore, we focused on only Megabase-scale TADs reported in an independent study by Ren and colleagues (see **Supplementary Methods**) in human ES cells (n=2,502) and ES cell-derived differentiated cells, including: mesendoderm (n=2,479), mesenchymal stem cells (n=2,290), neural progenitor cells (n=2,378) and trophoblast-like cells (n=2,435) ^18^. We observed that the majority of daSTRs at human ES cell TAD boundaries reported in Schmitt et al. were also observed at boundaries invariant across the other four ES cell-derived differentiated cell types (**Supplementary Fig. 7A-B**). Moreover, 14 of the 15 daSTRs associated with human ES cell TAD boundaries were also found at boundaries in mouse ES cells, indicating strong conservation across species (**Supplementary Fig. 8**). Noteworthy, nearly all daSTRs known to display paternal instability ^4,5^ showed co-localization to boundaries in mouse sperm and not in mouse oocytes ^19^, suggesting that daSTRs are at boundaries in the cell type and developmental timing when germ line expansion takes place (**Fig. 2, Supplementary Fig. 9**). Together, these results indicate that genomic loci susceptible to unstable repeat expansion are present at boundaries that exist in embryonic and somatic cell lineages and present in sperm when the instability is paternally inherited.

**Figure 2.**
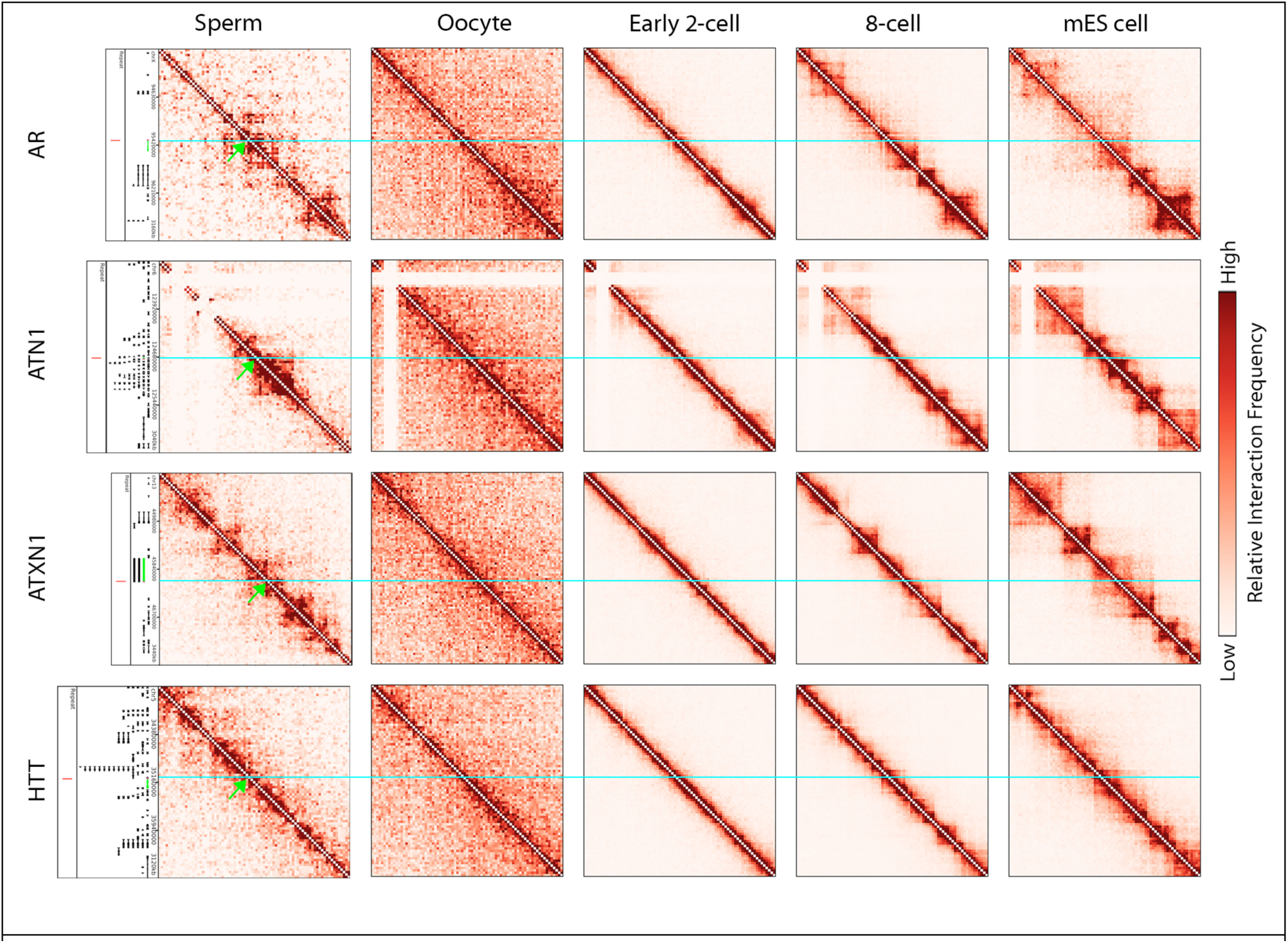
Repeats undergoing pathologic unstable expansion in the paternal germline are positioned at chromatin domain boundaries in mouse sperm. Hi-C heatmaps at repeat expansion loci across several stages in early murine development. Neon blue line indicates the genomic location of repeat tracts. Data analyzed from ^19^.

To understand genome folding in a tissue relevant to the subset of STR expansions linked to neurological dysfunction (**Supplementary Table 1**), we next analyzed recently published Hi-C data from human fetal cortical tissue ^17^. We applied the DI+HMM method to detect 2,102 TADs and a full sweep of 9,473 TAD/subTADs in human cortical plate tissue ^17^ (detailed in **Supplementary Methods**). We focused our analysis on the 22 daSTR loci specifically associated with neurological or neuromuscular disorders. Consistent with our observations in human ES cells, we found 13 out of the 22 neurological daSTR loci were detected at boundaries in human fetal cortical tissue (**Supplementary Figs. 10-11)**. An additional 7 daSTRs localized to qualitatively apparent boundaries (**Supplementary Figs. 12-13**). These results indicate that the large majority of the neurological daSTRs (20 out of 22) are located at domain boundaries in human cortical tissue (**Supplementary Fig. 14**). Thus, although we do not know the target cell type for many unstable repeat expansion disorders, the strong enrichment of daSTRs at boundaries regardless of tissue origin suggests that boundary location is robust to the cell types relevant for the pathology.

Due to the diversity of pathologic STR attributes (**Supplementary Table 1**), we sought to understand if a particular repeat class was driving the co-localization with boundaries. We stratified our daSTRs into 4 main groups: (i) a CAG repeat unit in exons or 5’UTR (n=9), (ii) a GCG repeat unit in exons or 5’UTR (n=8), (iii) repeat units in introns (n=5), and (iv) a CTG repeat unit in 3’UTRs (n=3) (**Table 2, Supplementary Methods**). Although statistical power was restricted by the small size of the groups, all four classes of daSTRs showed strong enrichment at boundaries compared to other normal-length repeats matched by sequence and gene body location (**Supplementary Fig. 15**). Together, these results indicate that a diverse range of loci linked to repeat expansion disorders are markedly enriched at domain boundaries even above normal-length repeats.

To understand the molecular mechanisms underlying the placement of daSTRs with respect to genome folding, we explored the genetic and epigenetic properties of domain boundaries. Consistent with previous reports, we found high enrichment for the architectural protein CCCTC-binding factor (CTCF) at human ES cell boundaries compared to loci internal to domains (**Supplementary Fig. 16A-B**) ^9–11^. Noteworthy, we also observed a marked increase in the density of CpG islands at boundaries vs. non-boundaries, whereas classic repressive chromatin marks such as H3K9me3 were slightly depleted at boundaries, as expected, compared to loci internal to domains. Two classes of boundaries emerge: boundaries with high CpG island density and boundaries depleted of CpG islands (**Supplementary Fig. 16C-D**). Although CTCF binds at both boundary and non-boundary regions in the genome, the boundaries with the highest occupancy of CTCF also contain high density of CpG islands. These results demonstrate that TAD/subTAD boundaries represent hotspots in the genome of ultra-high density of CTCF occupancy and CpG islands.

We hypothesized that CpG island-rich boundaries might be mechanistically linked to the susceptibility of some loci to undergo unstable repeat expansion. To test this hypothesis, we stratified boundaries into those with daSTRs, those with genome-wide normal-length matched repeat tracts, and those that do not contain repeats. Boundaries containing matched normal-length repeat tracts show a striking shift in CpG island density compared to boundaries without repeats (Odds Ratio = 4.28, Fisher’s Exact Test, P = 3.9E-96, **Figs. 3A-B**). Importantly, daSTRs localize with boundaries exhibiting ultra-high CpG island density (blue spheres, **Fig. 3A**). Moreover, boundaries with daSTRs exhibit a dramatic increase in CpG island density even over the rigorous null of boundaries containing matched, normal-length repeats (Odds Ratio = 13.7, Fisher’s Exact Test, P = 3.56E-6, **Figs. 3B**). CpG island density was significantly higher at boundaries with daSTRs compared to matched repeats (empirical P=0.03, **Fig. 3C**). All four classes of daSTRs co-localized with boundaries showing higher CpG island density than the matched, normal length repeats (**Supplementary Fig. 17**). These data uncover unique genetic and epigenetic features (high CpG island density, high CTCF occupancy) at chromatin boundaries where daSTRs become unstable leading to repeat expansion disorders.

**Figure 3.**
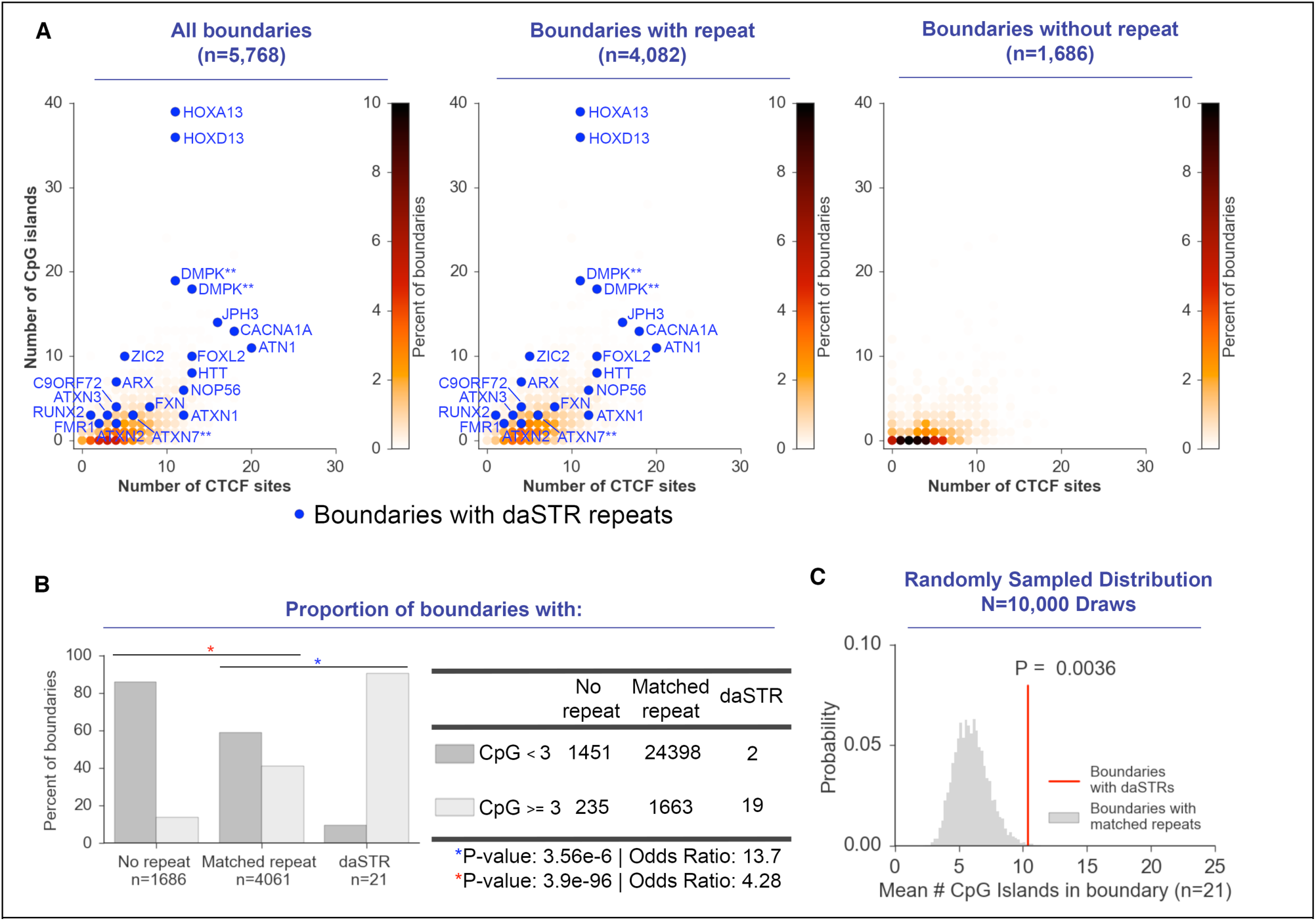
Boundaries containing disease-associated STRs (daSTRs) are characterized by ultra-high density of CpG islands. (A) Contour density plot depicting the number of CTCF sites versus CpG islands in 200kb bins representing boundaries with normal-length, matched repeats or those depleted of repeat tracts. Points are colored according to density. daSTRs are marked in blue. **(B)** 2x3 contingency table comparing CpG island density at boundaries with daSTRs, normal-length matched repeats, and no repeat tracts. **(C)** Randomization test comparing the number of CpG islands in boundaries with daSTRs or normal-length, matched repeats.

To investigate whether unstable, pathologic repeat expansion might disrupt higher-order chromatin structure at domain boundaries, we performed the Chromosome Conformation Capture Carbon Copy (5C) assay^20,21^ on B lymphocytes from a Fragile X Syndrome (FXS) patient with ∽935 CGG short tandem repeats and his healthy male sibling (GM09237 and GM09236 from the NIGMS Coriell Cell Repository) (**Figure 4A, Supp. Table 6**). We found a striking loss of intra-domain contacts and gain of cross-domain contacts at the *Fmr1* locus in the patient compared to the healthy sibling (**Figure 4B**). Notably, we did not observe any difference in boundary structure between patient and control when examining domains distal to *Fmr1* (**Figure 4C**), suggesting that loss of domain integrity in the FXS patient was specific to the repeat expansion. These results indicate that the *Fmr1* boundary is ablated in Fragile X Syndrome and open up new possibilities into understanding how mutation-length repeats perturb three-dimensional chromatin structure in other unstable repeat expansion diseases.

**Figure 4.**
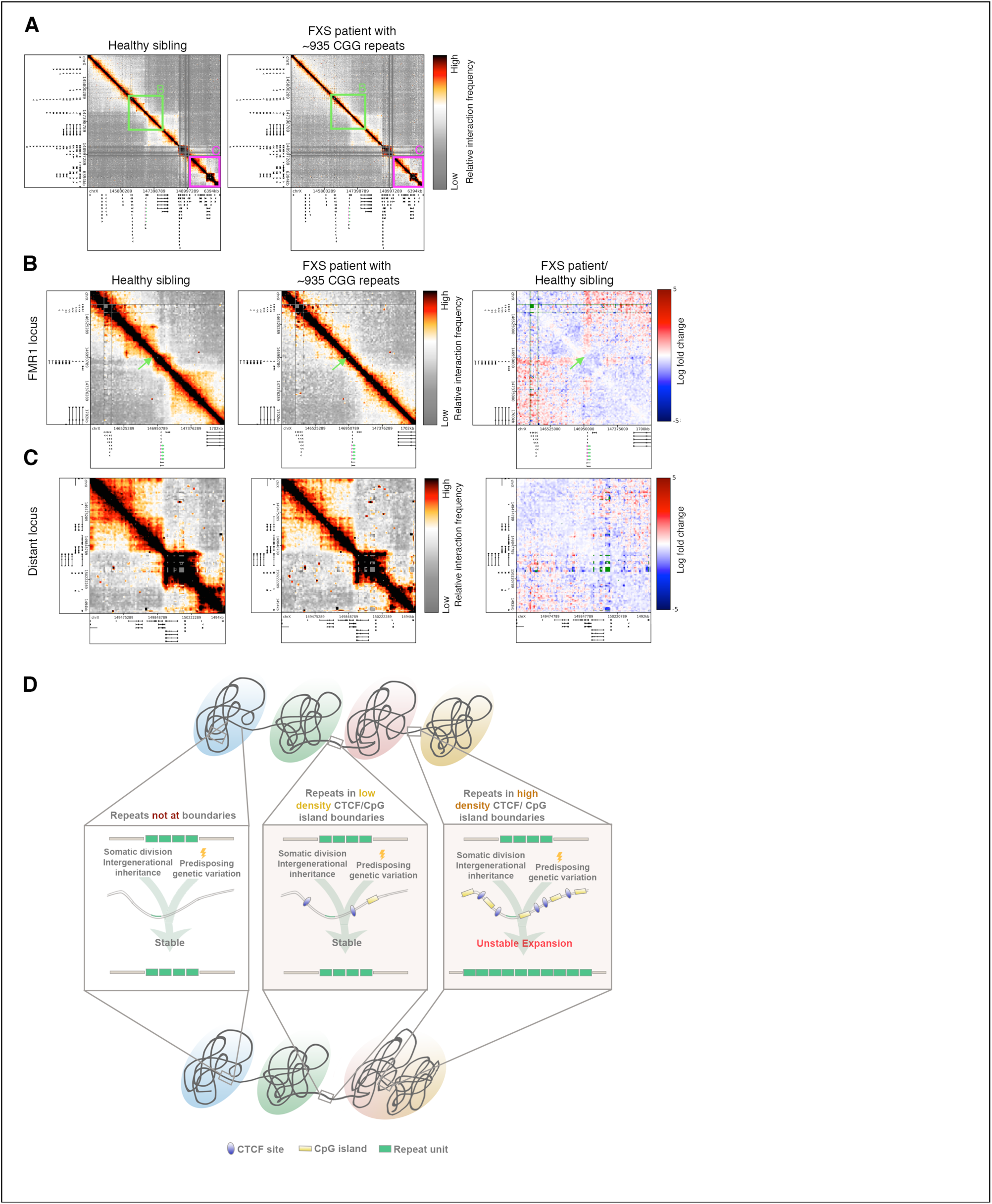
Unstable repeat expansion at the *Fmr1* gene is associated with domain boundary disruption in Fragile X Syndrome. (**A**) 5C contact matrices in B-lymphocytes from a male Fragile X Syndrome patient with ∽935 CGG repeats (Coriell Catalog ID GM09237) and a male healthy sibling (Coriell Catalog ID GM09236). The *Fmr1* gene is highlighted in green. **(B-C)** Zoom-ins on (**B**) the *Fmr1* locus and (**C**) a locus ∽3Mb downstream of *Fmr1* are shown for the Fragile X Syndrome patient and healthy sibling. The log fold change between the diseased and healthy sibling highlights contacts gained (red) and contacts depleted (blue). **(D)** Schematic for the role of high CpG island density at 3D genome folding domain boundaries on repeat tract instability in the human genome.

Genetic disruptions that alter chromatin domain integrity lead to ectopic, cross-boundary interactions and the disruption of gene expression ^13,14,22–25^. We reasoned that unstable daSTR expansion might lead to the loss of functional boundary insulation in patients with trinucleotide repeat expansion disorders. To test this idea, we analyzed data from two independent studies profiling genome-wide gene expression in human prefrontal cortex tissue from Huntington’s disease patients and healthy human controls ^26,27^ (**Supplementary Methods**). We found a decrease in insulation between TADs demarcating the repeat in the *HTT* gene, as evidenced by an increase in correlated expression of a cross-boundary gene pair (i.e. *GRK4* vs. *HTT*) in patients vs. controls (**Supplementary Figs. 18A-B, 19A**). By contrast, and consistent with previous reports ^13,25^, genes residing in the same domain (i.e. *GRK4* vs. *MFSD10*) did not show an alteration in expression correlation profiles in patients vs. controls (**Supplementary Figs. 18C, 19B**). These data indicate that mutation-length expansion at the *HTT* boundary results in loss of insulation between adjacent domains in Huntington’s disease and raise the intriguing possibility that boundaries might also be functionally disrupted in repeat expansion disorders.

Our data support a working model that sheds new light on the fundamental question of why key locations in the genome undergo unstable STR expansion, whereas tens of thousands of normal-length STR tracts across the genome remain stable. We demonstrate that although repeat expansion disease associated STR loci are diverse in the type, length and location of the repeat tract, they appear to share a common spatial placement at boundaries of 3D genome folding domains. We uncover that 3D genome folding domain boundaries are hotspots for high-density localization of CpG islands and CTCF sites, both of which are epigenetic features that have been linked to repeat instability ^28–31^. Studies in model organisms show that mutations in genes encoding key machinery involved in DNA replication, repair and recombination result in repeat expansion ^1,4,5,32^. In humans, a recent genome-wide association study in Huntington’s disease patients identified a link between genetic variation in DNA repair machinery and the age of onset of the disease^33^. Our study does not aim to address the genetic variation associated with individuals who get repeat expansion. Rather, we propose a working model in which 3D chromatin domain boundaries with high CpG island density are highly susceptible to unstable STR expansion in the case of predisposing genetic variation compared to any other location in the genome (**Fig. 4D**). There is a severe paucity of daSTRs at genomic loci internal to domains and boundaries without CpG islands.

Recent high-resolution Chromosome-Conformation-Capture sequencing studies have revealed that TAD boundaries are perturbed in rare human limb malformation diseases ^22^ and certain types of cancers ^23,25^, leading to enhancer miswiring and pathogenic disruption of domain integrity. An important prediction from our model is that boundaries might be disrupted in unstable repeat expansion disorders (**Fig. 4D**). We report the first evidence to our knowledge suggesting that TAD boundaries can be structurally and functionally disrupted in Fragile X Syndrome and Huntington’s disease. Altogether, our data reveal a fundamentally new link between higher-order 3D genome folding and trinucleotide repeat expansion disorders. An exciting area of future inquiry will be determining whether pathologic repeat instability causes or is caused by domain boundary disruption. Future studies unraveling the causal relationship between chromatin structure and repeat instability will illuminate the potential of topology-directed therapy in treating disease.

## Acknowledgements

Jennifer E. Phillips-Cremins is a New York Stem Cell Foundation – Robertson Investigator, an Alfred P. Sloan Foundation Fellow and a primary member of the Epigenetics Institute at the University of Pennsylvania. This research was supported by The New York Stem Cell Foundation (J.E.P.C), the Alfred P. Sloan Foundation (J.E.P.C), the NIH Director’s New Innovator Award from the National Institute of Mental Health (1DP2MH11024701; J.E.P.C), a 4D Nucleome Common Fund grant (1U01HL12999801; J.E.P.C) and a joint NSF-NIGMS grant to support research at the interface of the biological and mathematical sciences (1562665; J.E.P.C).

